# Environmental chemical diethylhexyl phthalate alters intestinal microbiota community structure and metabolite profile in mice

**DOI:** 10.1101/581975

**Authors:** Ming Lei, Rani Menon, Sara Manteiga, Nicholas Alden, Carrie Hunt, Robert C. Alaniz, Kyongbum Lee, Arul Jayaraman

**Affiliations:** Department of Chemical and Biological Engineering, Tufts University, Medford, MA 02155; Artie McFerrin Department of Chemical Engineering, Texas A&M University, College Station, Texas 77843; Department of Biomedical Engineering, Texas A&M University, College Station, TX 77843; Department of Microbial Pathogenesis and Immunology, College of Medicine, Texas Health Science Center, Texas A&M University, College Station, TX 77843; Artie McFerrin Department of Chemical Engineering, Texas A&M University, College Station, TX 77843

## Abstract

Exposure to environmental chemicals during windows of development is a potentially contributing factor in gut microbiota dysbiosis, and linked to chronic diseases and developmental disorders. We used a community-level model of microbiota metabolism to investigate the effects of diethylhexyl phthalate (DEHP), a ubiquitous plasticizer implicated in neurodevelopmental disorders, on the composition and metabolite outputs of gut microbiota in young mice. Administration of DEHP by oral gavage increased the abundance of *Lachnoclostridum*, while decreasing *Akkermansia, Odoribacter*, and *Clostridium sensu stricto*. Addition of DEHP to *in vitro* cultured cecal microbiota increased the abundance of *Alistipes, Paenibacillus*, and *Lachnoclostridium*. Untargeted metabolomics showed that DEHP broadly altered the metabolite profile in the culture. Notably, DEHP enhanced the production of *p*-cresol, while inhibiting butyrate synthesis. Metabolic model-guided correlation analysis indicated that the likely sources of *p*-cresol are *Clostridium* species. Our results suggest that DEHP can directly modify the microbiota to affect production of bacterial metabolites linked with neurodevelopmental disorders.

**Importance:** Several previous studies have pointed to environmental chemical exposure during windows of development as a contributing factor in neurodevelopmental disorders, and correlated these disorders with microbiota dysbiosis, little is known about how the chemicals specifically alter the microbiota to interfere with development. The findings reported in this paper unambiguously establish that a pollutant linked with neurodevelopmental disorders can directly modify the microbiota to promote the production of a potentially toxic metabolite (*p*-cresol) that has also been correlated with neurodevelopmental disorders. Further, we use a novel modeling strategy to identify the responsible enzymes and bacterial sources of this metabolite. To the best of our knowledge, the present study is the first to characterize the functional consequence of phthalate exposure on a developed microbiota. Our results suggest that specific bacterial pathways could be developed as diagnostic and therapeutic targets against health risks posed by ingestion of environmental chemicals.

## Introduction

The mammalian gastrointestinal (GI) tract harbors microbial communities that impact a wide array of physiological functions, including digestion, immune system development, and defense against pathogens. Alterations in the microbiota composition leading to functional imbalance, or dysbiosis, have been linked to various chronic diseases and disorders, including inflammatory bowel disease (1), colorectal cancer (2), fatty liver disease (3), diabetes (4), and neurodevelopmental disorders (5). Factors known to cause dysbiosis include diet, infection, and use of antibiotics. In recent years, environmental chemicals have emerged as another factor contributing to alterations in the microbiota.

Exposure to biologically active synthetic chemicals present in household and industrial products, particularly during critical windows of development, has been shown to result in microbiota dysbiosis and correlated to various disorders of the immune and nervous systems (6). An environmental chemical that is pervasive in the environment due to its widespread use as a plasticizer is diethylhexyl phthalate (DEHP) (7). In vertebrate animals, DEHP impacts reproduction and development (8). A recent study found increased serum DEHP concentrations in children diagnosed with autism spectrum disorder (ASD) (9). Additionally, fecal samples from children diagnosed with ASD have elevated concentrations of bacterial metabolites such as *p*-cresol (10), pointing to a potential link between the health effects of DEHP exposure and the intestinal microbiota.

This link is supported by multiple studies with other environmental chemicals correlating microbiota dysbiosis with adverse effects of exposure. A study in mice showed that exposure to benzo[a]pyrene resulted in pronounced alterations of the intestinal microbiota, including a decrease in the abundance of *Akkermansia muciniphila*, and an increase in the levels of inflammatory indicators (11). Another recent study found that Bisphenol-A (BPA) exposure exacerbated the effects of chemically-induced colitis, and that these effects were accompanied by altered fecal levels of tryptophan-derived metabolites (12).

The importance of bacterially produced metabolites in dysbiosis-related disorders was highlighted by Hsiao et al., who showed that the behavioral abnormalities observed in a maternal immune activation (MIA) model of anxiety-like behavior in mice correlated with changes in the abundance of intestinal bacteria and the concentration of bacterial metabolites in serum (13). This study also showed that the behavioral abnormalities in the MIA model could be improved by controlling the level of a specific tyrosine metabolite, 4-ethylphenysulfate. Altered levels of microbiota-associated metabolites such as indoxyl sulfate and *p*-hydroxyphenyllactate have also been detected in blood and urine of children diagnosed with ASD (14, 15), suggesting that the link between bacterial metabolites and neurodevelopmental disorders could be relevant in humans. On the other hand, it is unclear whether the aforementioned alterations in bacterial metabolite profiles directly result from environmental chemical exposure. While several studies have investigated the effects of DEHP exposure on host reproductive, nervous and metabolic tissues (16–18), little is known about the impact of this ubiquitous chemical on intestinal microbiota composition and function.

A majority of studies have used early life exposure models to study the effects of environmental chemicals on the intestinal microbiota. Observations from these studies suggest that changes to the intestinal microbiota can persist beyond the pre- or perinatal exposure period (19, 20). For example, perinatal exposure of rabbits to BPA *in utero* and during the first week of nursing led to a decrease in short-chain fatty acid (SCFA) producing bacteria at 6 weeks of age (21). Another study found that continuous exposure to diethyl phthalate, methylparaben, and triclosan beginning at birth induced significant changes to the gut microbiota of adolescent rats (22). To-date, few studies have looked at the effect of environmental chemical exposure on a developed microbiota.

While *in vivo* studies on the microbiota offer physiologically relevant insights, they can be often difficult to interpret due to confounding influences from the host (23). Apart from directly altering microbiota composition, environmental chemicals can also indirectly cause microbiota dysbiosis, for example by bringing about intestinal inflammation through activation of host receptors (24). Moreover, receptor activation can occur through an intestinal or liver biotransformation product, rather than the chemical itself (25). To elucidate the mechanistic role of dysbiosis resulting from environmental chemical exposure in a particular disease or disorder, it is important to delineate the effects of environmental chemical exposure on the intestinal microbiota from those on the host.

In this work, we used *in vivo* exposure in mice and an *in vitro* culture model to investigate the effect of DEHP on the intestinal microbiota composition and its metabolite output. Our results suggest that environmental chemical exposure can directly modify the intestinal microbiota to increase production of potentially neurotoxic microbial metabolites linked with behavioral abnormalities.

## Results

### Gut microbiota composition is altered in vivo in a time dependent manner

We investigated the effect of DEHP exposure on the gut microbiota by administering the chemical to 6-8 week old female C57/BL6 mice via oral gavage and analyzing the changes in the fecal microbial community at day 7 and day 14 post-exposure using 16S rRNA sequencing. Principal component analysis (PCA) of operational taxonomic unit (OTU) counts showed samples grouping together by time point but not by DEHP treatment (Figure 1A), suggesting that changes in OTU profile driven by the host’s age may dominate over DEHP-driven changes. This trend agreed with classification results from partial least squares-discriminant analysis (PLS-DA), which achieved stronger separation between OTU profiles when samples were classified based on time point than chemical treatment (Figure S1). The results from PLS-DA also indicated that the effect of DEHP on the OTU profiles was greater on day 7 than day 14. Consistent with this observation, samples from DEHP treated mice showed a higher alpha diversity (Chao1 index) than control mice on day 7, but not day 14 (Figure 1B). Similarly, linear discriminant analysis of the effect size (LefSe) identified a larger number of OTUs affected by DEHP treatment in day 7 samples compared to day 14 samples (Figure 1C). At the genus level, the abundance of *Akkermansia, Odoribacter*, and *Clostridium sensu stricto* decreased in the DEHP samples collected on day 7. Of these, only *Clostridium sensu stricto* remained significantly depleted in the day 14 samples. An unclassified genus belonging to the order *Mollicutes RF9* was increased in abundance in the DEHP samples on day 7, and *Lachnoclostridum* was increased on day 14.

**Figure 1.**
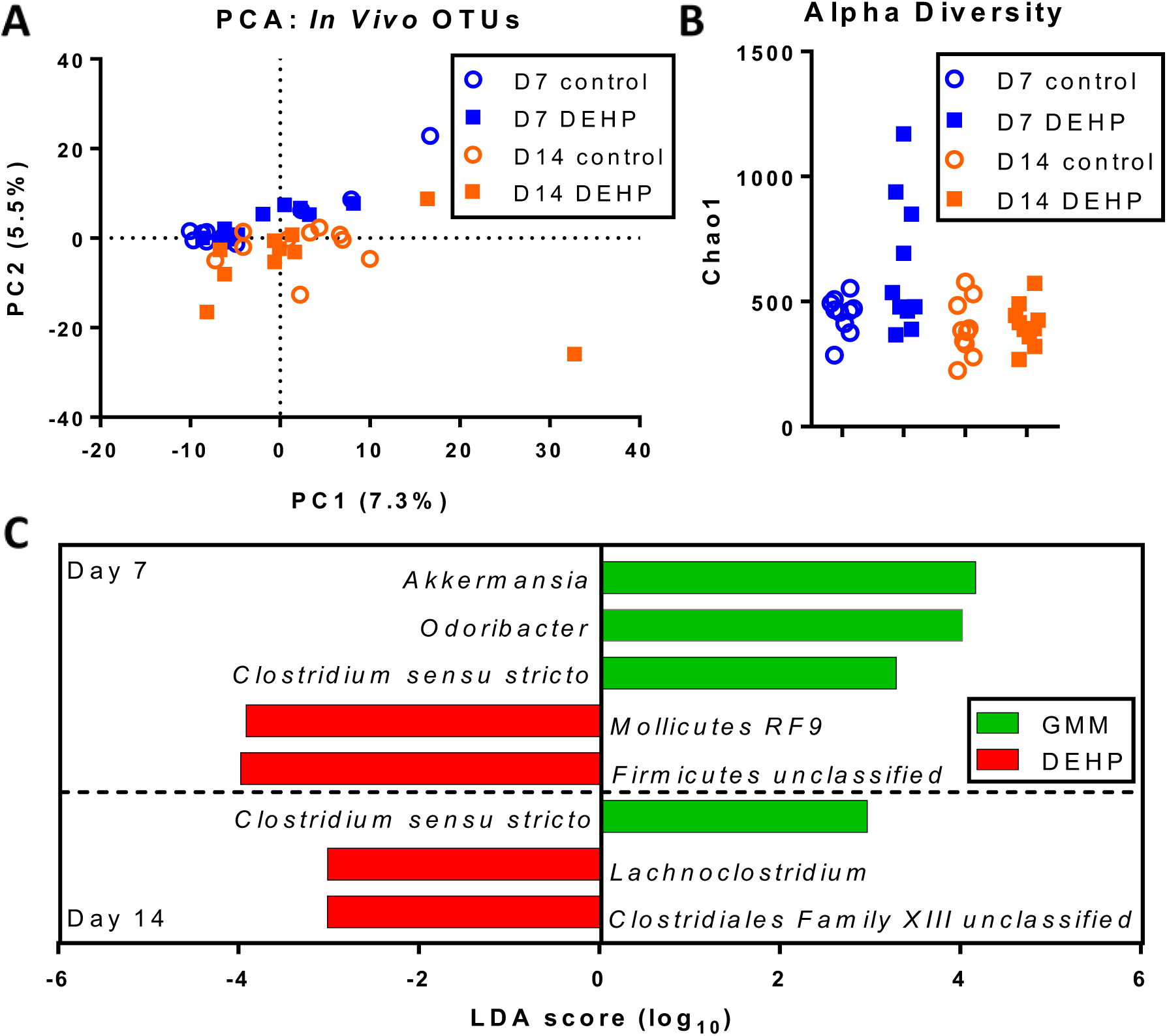
Metagenomic (16S rRNA) analysis of fecal microbiota from DEHP-exposed mice. (A) PCA on OTU counts. The percentage represents the percent variance explained by each axis. (B) Alpha diversity and (C) LefSE analysis of fecal microbiota OTU counts.

### Fecal metabolite profile is more strongly influenced by host-dependent factors than DEHP treatment

To determine whether the phthalate exposure also altered the profile of intestinal metabolites, we analyzed the fecal material using untargeted LC-MS metabolomics. We first confirmed that the orally administered DEHP was available to the microbiota by identifying the presence of mono(2-ethylhexyl)phthalate (MEHP), a product of enzyme-catalyzed DEHP degradation (26) in the metabolite data. As expected, MEHP was detected in fecal samples from DEHP treated mice, but not control mice (Figure 2A). Similar to the OTU profiles, results of PCA on the LC-MS features indicated that DEHP had a lesser effect on the fecal metabolite profile than the host’s age (Figure 2B). This result was consistent with two-tailed t-tests performed on individual data features, i.e., metabolites, which found very few statistically significant differences (none that could be assigned a putative identity) between time-matched samples from control and DEHP-treated mice on days 0, 7, and 14 (less than 1.6, 2.3, and 8.0% of total detected features, respectively). We detected a larger number of features that were significantly elevated or reduced in the day 14 fecal samples relative to day 0 and 7 samples (10.9 and 23.7%, respectively). These trends suggested that the global profile of fecal metabolites is more strongly influenced by host factors such as aging. To more directly assess the effects of DEHP on the microbiota in isolation from host influences, we performed the DEHP exposure experiment using an anaerobic batch culture model of murine cecal microbiota.

**Figure 2.**
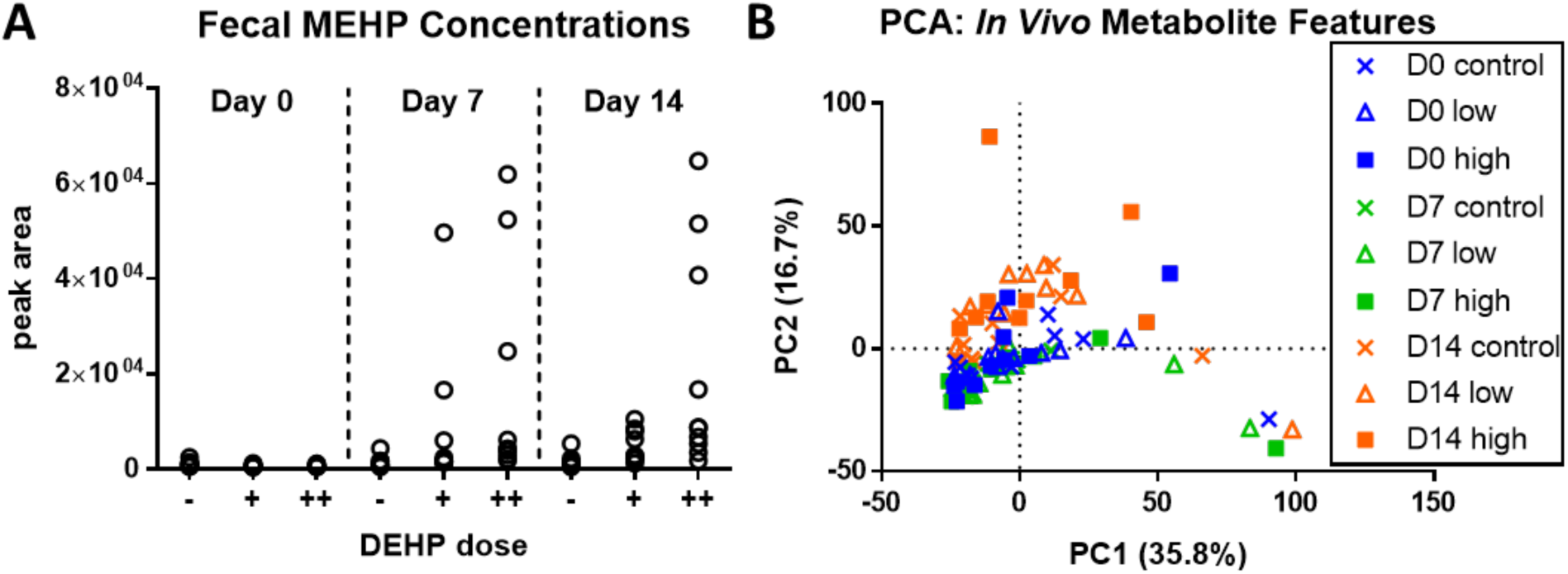
Metabolite analysis of fecal microbiota from DEHP-exposed mice. (A) LC-MS identification of MEHP in fecal material collected at day 7 and 14 from animals fed a low (+) or high dose (++) of DEHP. The level of MEHP in control samples (-) was below the limit of detection. (B) Scatter plot of the first two PC scores from PCA of the metabolite data.

### Anaerobic cecal batch culture captures in vivo microbiota diversity

We assessed whether the cecal contents culture could represent the biochemical diversity of murine cecal microbiota by characterizing the OTU profile of control cultures in gut microbiota medium (GMM) (27) without DEHP. Using QIIME with the SILVA database (28) as the reference, we identified approximately 2,000 distinct OTUs that are present on both days 1 and 7 of the culture. Nearly all of the OTUs (99.4 %) belong to four bacterial phyla: *Actinobacteria, Bacteroidetes, Firmicutes*, and *Proteobacteria* (Figure 3A). The number of families and genera represented in the culture were 65 and 119, respectively. A comparison of OTUs detected on day 7 of cultured cecal contents with cecal contents harvested from 8-week old female C56/BL6 mice showed 100% similarity in relative abundance at the phylum level and greater than 70% similarity at the genus and species levels (Figure S2). From day 1 to 7, there were significant shifts in the relative abundances of the OTUs. At the genus level, *Lactobacillus* and *Parabacteroides*, respectively, showed the maximum decrease and increase in terms of OTU counts, whereas *Fluviicola* and *Enterococcus*, respectively, showed the greatest decrease and increase in terms of fold-change (Figure 3B).

**Figure 3.**
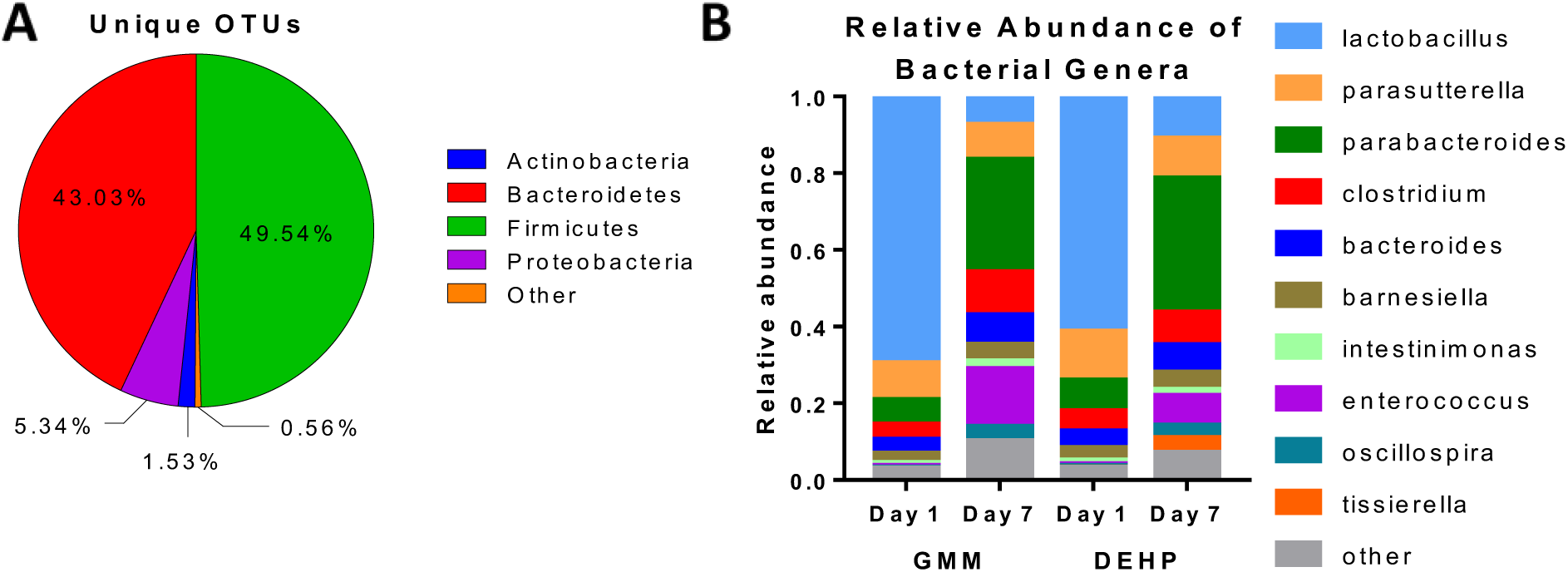
Metagenomic analysis of *in vitro* cecal luminal contents culture. (A) Phylum-level classification of unique OTUs in DEHP-treated cultures. (B) Relative abundance of bacterial genera on day 1 and 7 in control (GMM) and DEHP-treated cultures.

### Cecal culture produces a diverse array of secondary and amino acid metabolites

We next characterized the major metabolic products and substrates of the cecal cultures using untargeted LC-MS experiments. Principal component analysis (PCA) performed on the untargeted LC-MS features showed clear separation between day 1 and 7 samples from the inoculated GMM cultures (Figure 4A). In contrast, day 1 and 7 samples from culture tubes containing GMM without cells grouped closely together. Hierarchical clustering of the LC-MS data features revealed five distinct patterns (Figure 4B). The first group of features represented metabolites that were rapidly consumed and significantly depleted by day 1. A second, larger group was consumed more slowly, with significant depletion occurring only by day 7. The third group comprised rapidly produced metabolites that were significantly elevated by day 1. The fourth group of metabolites was produced more slowly, and significantly elevated by day 7, but not day 1. The fifth group was significantly elevated by day 1, but reduced by day 7 (Figure 4C).

**Figure 4.**
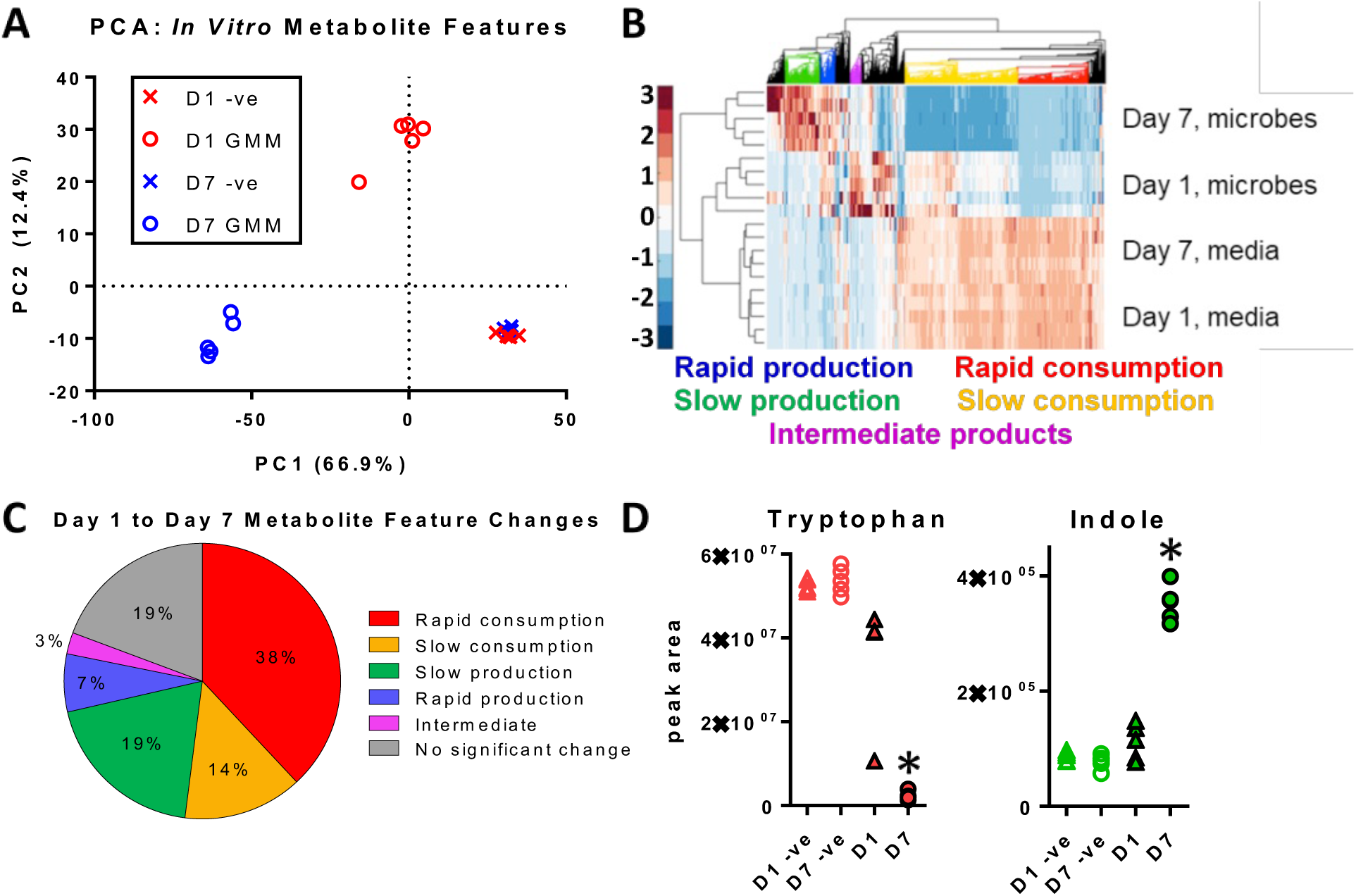
Metabolite profiles from *in vitro* cultured cecal luminal contents. (A) Scatter plot of the first two scores from PCA representing microbial metabolites produced on day 1 and day 7. (B) Heat map of detected ion peaks with different patterns of substrate utilization and product formation. (C) Percentage distribution of detected features classified as products, substrates, or intermediates based on their time profiles. (D) Profiles of tryptophan and indole in the cecal luminal content cultures. Filled and open markers represent inoculated cultures and tubes incubated without luminal contents, respectively. Triangles and circles represent day 1 and 7 time points, respectively. The colors correspond to the classifications in the heat map and pie chart. *: *p*-value<0.05 when compared to inoculated culture at day 1 (two-tailed t-test).

Putative metabolite identities were assigned to the LC-MS data features based on accurate mass and product ion spectra (MS/MS data). Overall, 118 and 156 of the features in the positive and negative mode ionization data, respectively, were assigned a putative KEGG compound identifier. The merged list of putatively identified metabolites from both ionization modes comprised 204 unique compounds (Table S1). The overlap in metabolites identified by positive and negative mode experiments was very small (27/231), indicating that using two different LC-MS methods significantly broadened coverage.

We utilized the Search Pathway tool of KEGG Mapper to associate each putatively identified metabolite with one or more functional categories (Table S2). Based on this mapping, the largest KEGG function categories were biosynthesis of secondary metabolites (54 mapped metabolites), microbial metabolism in diverse environments (47), and biosynthesis of antibiotics (35). Additional pathways captured by the data included fermentative reactions known to occur in the intestine, such as L-carnitine metabolism to gamma-butyrobetaine and trimethylamine. If the different amino acid metabolism subcategories were pooled, then more than one-third (79/204) of the metabolites belonged to this more general category. Interestingly, we detected the production of the neurotransmitter serotonin, despite the absence of any host cells in the cultures. Other aromatic amino acid (AAA) products including the tryptophan metabolites indole (Figure 4D), indole-3-propionic acid and indole-3-carboxylic acid were also detected. Phenylalanine-derived metabolites detected in the cultures included phenylacetic acid, phenylpropionic acid, and 3-(3-hydroxyphenyl)propionic acid, and tyrosine-derived metabolites detected included *p*-cresol and *p*-hydroxyphenylacetic acid.

### Metabolic function in cecal cultures is distributed heterogeneously across taxonomic groups

We next analyzed the genomes of bacterial groups detected in the culture to characterize the enzymatic reactions responsible for the metabolic products. Cross-referencing the list of identified OTUs against KEGG and UniProt, we obtained a model that included at least one annotated genome for 94 of 119 genera detected in the cultures, accounting for greater than 96% of the bacterial counts (Figure 5A).

**Figure 5.**
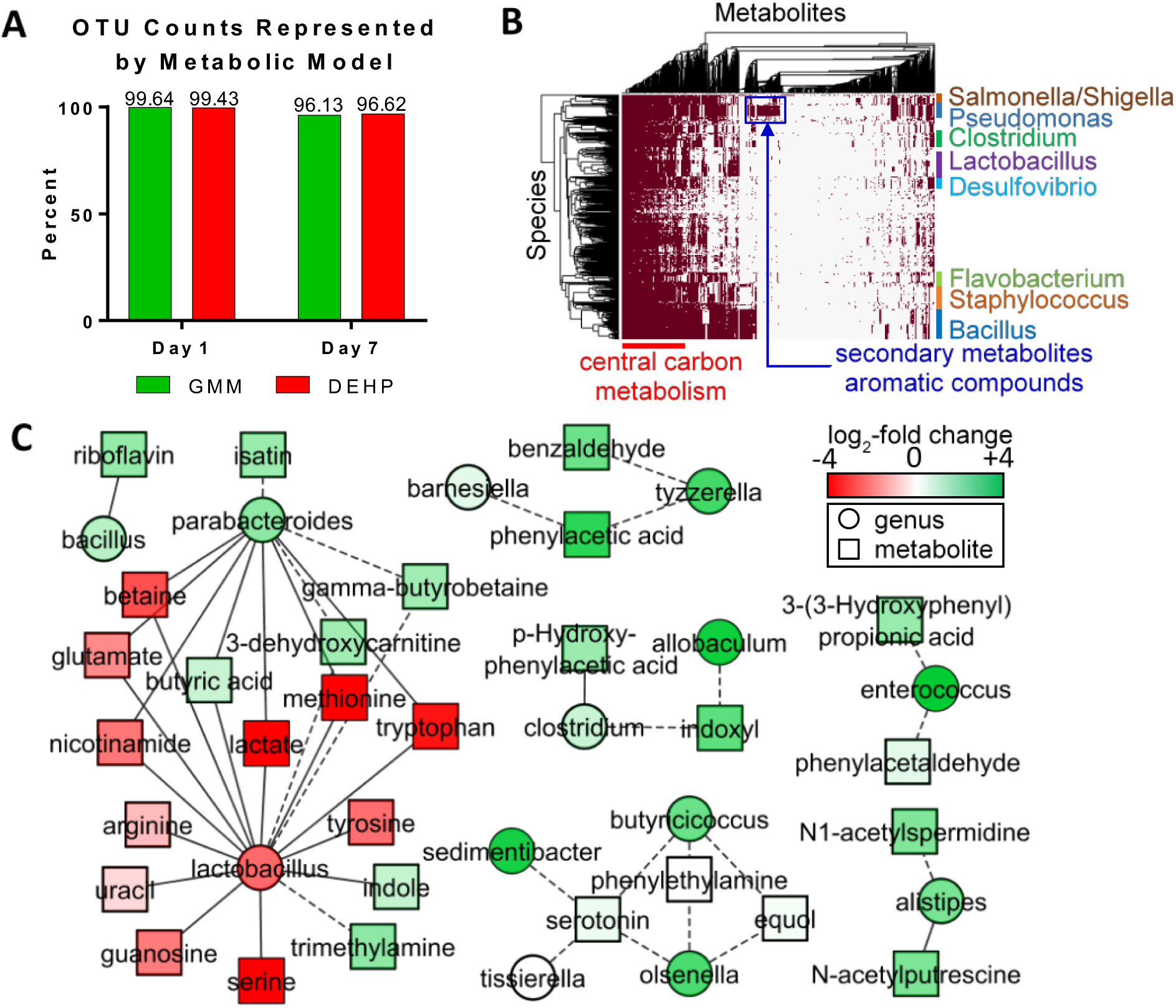
Model of metabolic reactions in *in vitro* culture of cecal luminal contents. (A) Fraction of genus-level OTU counts represented by the metabolic model. (B) Hierarchical clustering of genera and metabolites in the model. (C) Correlation network showing significant Pearson correlations between genera (circles) and metabolites (squares). Fold-change from day 1 to 7 is indicated by red (decrease) and green (increase) colors. Solid edges between nodes indicate that the genus has at least one species capable of metabolizing the connected metabolite (per database annotation of the genome), while dotted lines indicate a purely empirical correlation.

We evaluated the coverage of metabolic functions represented in the model by comparing the orthologs of the model against the orthologs predicted by Tax4Fun (29), using 16S sequence data, reference sequences in the SILVA database, and annotated genomes cataloged in KEGG as inputs. Based on KEGG orthology (KO) numbers, 83% of the gene functions predicted by Tax4Fun overlapped with the estimates from our model. When weighted by the relative abundance of the OTUs, we found a strong correlation between ortholog counts from Tax4Fun and our model (Figure S3). Despite the similarity in functional coverage with our model, the Tax4Fun prediction included many additional organisms most of which (more than 80%) did not match the OTUs classified by the SILVA analysis on the cecal culture. Thus, our model parsimoniously covered the biochemical diversity of the cecal culture without overpredicting the underlying taxonomic diversity.

The molecular functions represented by KO numbers were used to associate the bacterial species in our model with enzymatic reactions and their corresponding metabolites. The reactions distributed highly unevenly across the different genera (Figure 5B). Out of more than 4,000 reactions, only 4 (involved in DNA replication) mapped to every genus. Approximately 9% (349/4034) of the reactions mapped to a single genus, indicating that certain metabolic functions (e.g., carotenoid biosynthesis, steroid hormone biosynthesis, methane metabolism, glycosphingolipid biosynthesis, benzoate degradation) required the participation of particular genera. Based on reaction definitions in KEGG, 84% (49/58) of the confidently identified metabolites (Table S3) were mapped to one or more organisms in the model. Similar to the trend for reactions in the model, the metabolites also distributed heterogeneously across the different organisms. Amino acids (e.g., phenylalanine, tryptophan), nucleosides/nucleotides (e.g., uridine, uracil), and vitamins (e.g., riboflavin) associated with nearly ubiquitous reactions found in more than 90% of the genera, whereas certain fermentation products (e.g., benzaldehyde, indole-3-acetate) associated with rare reactions found in less than 50% of the genera.

### Correlation analysis finds significant associations between metabolite levels and relative abundance of organisms in the culture

To identify significant associations between individual OTUs and metabolites, we calculated Pearson correlation coefficients (PCCs) between the peak area of each confidently identified metabolite and the relative abundance of each highly abundant genus (>0.05% of total OTU counts) detected on both days 1 and 7. After correcting for false discovery rate (FDR), we found 46 significant correlations. We mapped these correlations to the metabolic model described above to highlight different interactions between organisms and metabolites based on whether or not an organism possessed the enzyme to directly act on the correlated metabolite. The resulting correlation network is shown as connected nodes in a bipartite graph (Figure 5C). Excluding common amino acids and GMM components, the strongest correlations were: *Lactobacillus*-lactate (PCC=0.98), *Butyricicoccus*-serotonin (PCC=0.93) *Enterococcus*-3-(3-hydroxyphenyl) propionic acid (PCC=0.92), *Clostridium*-indoxyl (PCC=0.91), and *Alistipes*-N1-acetylspermidine (PCC=0.88). (Figure 5C, Table S4).

### DEHP induces changes in microbiota composition in vitro

We next examined the effects of DEHP on bacterial abundance in the cecal culture. At the genus level, DEHP increased the abundance of *Alistipes, Paenibacillus,* and *Lachnoclostridium* on day 1, while decreasing the abundance of *Fluviicola* and *Symbiobacterium*. On day 7, we detected an increase in *Tissierella* (Figure 6A). At the OTU level, DEHP increased *Alistipes putredinis, Lachnoclostridium bolteae,* and *Lachnoclostridium sacharolyticum* on day 1. On day 7, we detected an increase in *Tissierella praecuta*, and decreases in *Bacillus velezensis* and *Lactobacillus brevis*. Overall, DEHP exerted a greater effect on microbiota composition on day 1 compared to day 7, both in terms of number of altered OTUs as well as the changes in relative abundance of these OTUs.

**Figure 6.**
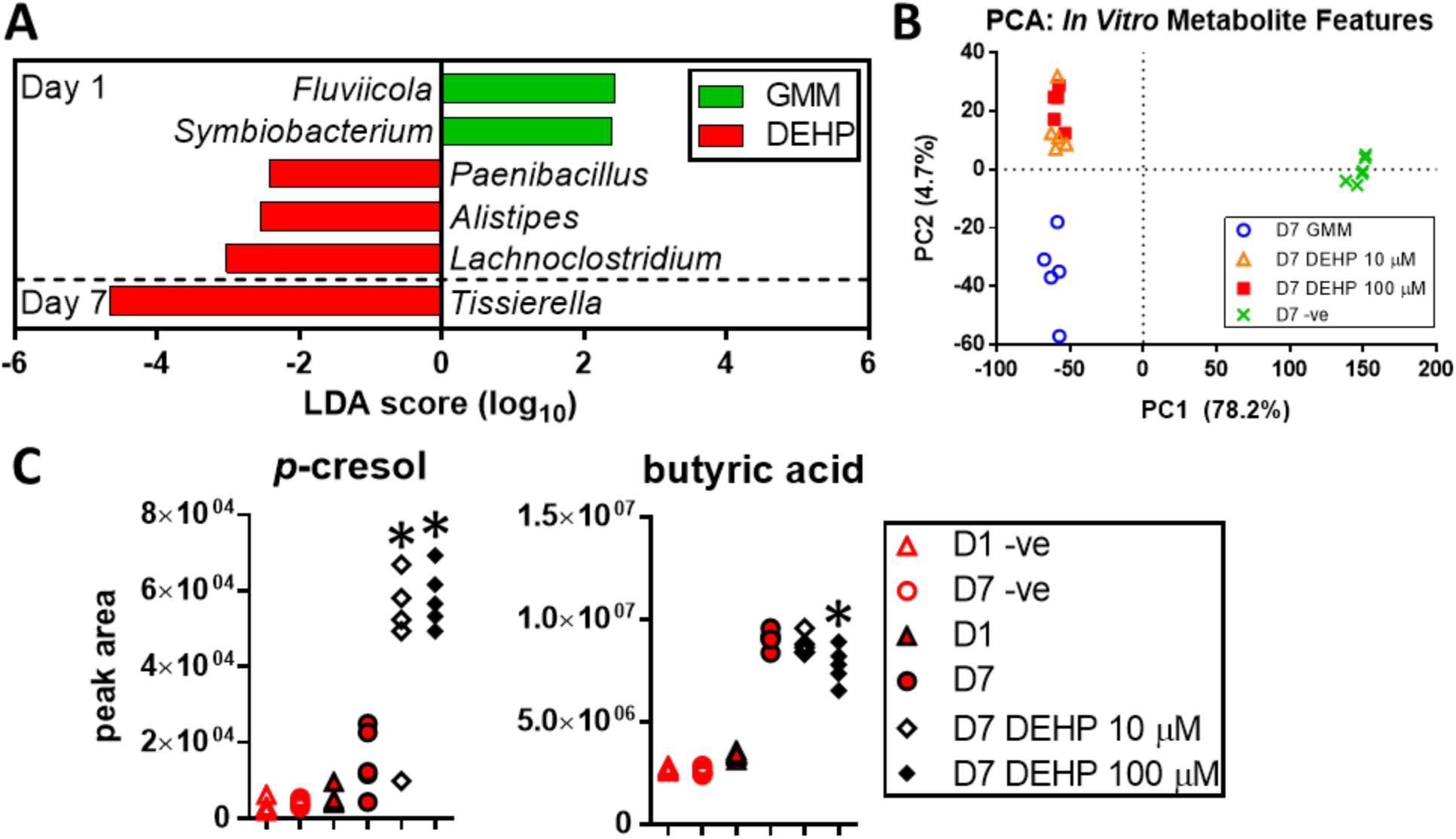
Significant microbial and metabolite changes in *in vitro* cultured cecal luminal contents with DEHP. (A) LefSe analysis of genus-level microbiota changes induced by DEHP. (B) Scatter plot of first two PC scores from PCA of metabolite features detected in the cecal cultures (positive mode IDA data). (C) Dose-dependent changes in *p*-cresol and butyric acid with DEHP on day 7. **\***: -value<0.05 when compared to day 7 culture without DEHP addition (two-tailed t-test).

### DEHP broadly alters metabolite profile of cecal microbiota

Addition of DEHP significantly altered the profile of metabolites in the cultures (Figure 4A). At 10 and 100 μ M, DEHP altered 16.8% and 20% of the LC-MS features detected on day 7 in the positive ionization mode experiment (Figure S4). The negative mode data showed even greater effects, with 46.7 and 47.2% of the detected features altered at 10 and 100 μ M, respectively. Only a subset of these features could be assigned a chemical identity due to lack of matching entries in reference databases. Similar to the in vivo exposure, we detected a dose-dependent accumulation of MEHP, a degradation product of DEHP (Figure S5), confirming that the cecal culture is also capable of this reaction. Figure 4C shows representative profiles of confirmed metabolic products that were increased (cresol) and decreased (butyric acid) by DEHP treatment. *p*-cresol was identified by matching both the m/z ratio and retention time. We observed a retention time shift running the LC method for samples vs pure standards, possibly due to matrix effects of the samples. To account for this shift, we used local linear regression (Figure S6). Additionally, oleic acid and linoleic acid were decreased in the DEHP-treated cultures on day 7, while isatin was increased. An additional feature detected in positive mode ionization at *m/z* 138.0886 had a dose-dependent increase in response to DEHP, and was putatively identified as tyramine, 1-methylnicotinamide, or 2-hydroxyphenethylamine. It was not possible to assign a unique identity, because the product ion spectrum indicated that the corresponding ion chromatogram peak could represent more than one compound.

## Discussion

Studies have implicated dysbiosis of the gut microbiota in developmental disorders associated with exposure to environmental toxicants (22, 30). In this work, we focus on the effects of DEHP, a pervasive environmental chemical and endocrine disruptor associated with neurodevelopmental disorders such as ASD (9). Previous *in vivo* studies (30, 31) used early-life exposure models to investigate the microbiota’s role in developmental health and disease. Fewer studies have investigated the effect of environmental chemical exposure on a developed microbial community in mammals. In the present study, we mimicked human exposure during adolescence by continuously exposing mice to DEHP from ages 6 to 8 weeks. Additionally, we used an anaerobic batch culture model to investigate the effects of DEHP on the microbiota community structure and metabolites.

Continuous DEHP exposure modestly increased the alpha-diversity of the fecal microbiota after 7 days, an effect that dissipated by day 14. While it is often assumed that a diverse gut microbiome is beneficial for the host, this is not necessarily the case, as gut health is also impacted by the microbial enterotype. After 14 days, mice exposed to DEHP showed an increased abundance of *Lachnoclostridum* and an unclassified genus of *Clostridiales Family* XIII. While the effect of DEHP on gut microbiota of young mice has not been previously reported, studies in humans have associated overrepresentation of *Lachnoclostridium* species with neurodevelopmental disorders such as ASD (32, 33). This suggests that the enterotype resulting from pollutant exposure could play a role in dysbiosis associated neurodevelopmental disorders.

We detected minimal changes in metabolite profile of fecal material from mice exposed to DEHP for 7 or 14 days. This could be due to several factors. First, host-driven changes, e.g., due to development and age, could mask subtler effects of phthalate exposure. Second, microbial metabolites can be taken up and transformed by the host, which limits the extent to which fecal metabolite analysis can capture the profile of microbiota metabolites in the intestine. Moreover, fecal metabolites comprise not only bacterial products, but also dietary residues and endogenous metabolites produced by the host. To address these issues, we investigated the effect of DEHP exposure on gut microbiota community structure and biochemical function using an *in vitro* model.

*In vitro* models of the intestinal microbiota vary in their complexity, depending on the intestinal location is being mimicked and the degree of biophysical detail being incorporated (34). In the present study, we used a relatively simple anaerobic batch culture model, as the focus was on capturing biochemical function. This model recapitulated up to 70% of the microbiota found in murine cecum at the genus level, including strict anaerobes. Importantly, the culture supported the production of metabolites typically associated with fermentation of sugar and amino acid residues by intestinal bacteria. As the culture system lacks host cells, all of the accumulating products are unequivocally sourced from the detected bacteria. Among the detected products are potentially toxic compounds derived from AAAs, e.g., phenylethylamine (35) and phenylacetic acid (36). We also detected metabolites that are normally present at low levels in the intestine of healthy individuals but are elevated in developmental disorders. For example, 3-phenylpropionic acid and 3-(3-hydroxyphenyl) propionic acid are precursors of 3-(3-hydroxyphenyl)-3 hydroxypropionic acid (HPHPA), a compound elevated in the urine of children diagnosed with ASD (37). Another useful feature of the culture model is that the correlations identified between different metabolites as well as metabolites and organisms can reveal the sources of particular metabolic products. For example, lactic acid accumulated in the culture by day 1 and was rapidly consumed by day 7, which correlated with a significant depletion in *Lactobacillus*, a known lactic acid producer. The depletion in lactic acid was significantly correlated with butyric acid production, consistent with previous reports on bacterial conversion of lactic acid to butyric acid *in vitro* (38). Taken together, these findings suggest that the anaerobic batch culture broadly captures representative metabolic functions of the murine gut microbiota, while facilitating identification of bacterial metabolites and their source organisms.

We further analyzed the organism-metabolite correlations using a metabolic model to associate the correlations with enzymatic pathways encoded in the genomes of detected OTUs. Several of these associations confirm previously reported findings. For example, our analysis links *Bacillus* with riboflavin, consistent with previous reports on the synthesis of this vitamin by intestinal *Bacillus* species (39). Likewise, members of the genus *Alistipes* possess the enzyme putrescine N-acetyltransferase (EC# 2.3.1.57), which converts putrescine into N-acetylputrescine (40). A third example is the production of *p*-cresol, which previous reports have attributed to *Clostridium* species (41). Our analysis links *Clostridium* to *p*-hydroxyphenylacetic acid, which is converted to *p*-cresol via *p*-hydoxyphenylacetate decarboxylase (42).

Not all of the significant correlations mapped to an enzyme in the metabolic model. One limitation of the model is that genome annotations in KEGG and UniProt are incomplete. For example, the annotations for *Parabacteroides* in the databases did not include an enzyme for butyric acid synthesis, but a recent study found *buk* (butyrate kinase) and *ptb* (phosphotransbutyrylase) in the genomes of intestinal *Parabacteroides* species (43). In this regard, correlations that do not map to a cataloged enzyme could facilitate discovery of previously unknown metabolic functions of intestinal bacteria. Two putatively identified compounds, indoxyl or oxindole and isatin, strongly correlated with the expansion of *Clostridium* and *Parabacteroides*, respectively. Currently known metabolic reactions that produce indoxyl and isatin are catalyzed by monooxygenases requiring oxygen (44). As evidenced by the growth of obligate anaerobes, molecular oxygen is absent in the cultures, suggesting that there could be alternative mechanisms of incorporating hydroxyl groups into metabolites. Indeed, oxindole and isatin have been reported to be products of anaerobic indole degradation (45, 46).

Gut microbes have an extensive capacity to break down xenobiotics, including environmental chemicals, which can modulate their toxicity and bioavailability in the host (47). As was the case *in vivo*, we detected a dose-dependent accumulation of MEHP in the cecal contents culture, indicating that the organisms expressing the required esterase are also present *in vitro.* Compared to *in vivo* exposure, we detected fewer changes in the microbiota composition upon DEHP addition to the culture medium. This is possibly due to the rapid degradation of DEHP, which was continuously administered to the mice, but added as a bolus at the start of the culture. Nevertheless, we observed features common to both *in vivo* and *in vitro* exposures. Similar to the *in vivo* experiment, we detected an increase in *Lachnoclostridium,* although this increase was transient in the cultures. Additionally, we detected a transient increase in *Alistipes*, which has also been reported for subjects diagnosed with ASD and related GI conditions (48).

Treatment with DEHP significantly altered the profile of metabolic products accumulating in the culture. Notably, DEHP increased the accumulation of *p*-cresol, a putative biomarker of ASD (49), while decreasing the levels of butyric acid, a bacterial metabolite benefiting intestinal immune homeostasis and offering neuroprotective effects (50). The likely source of *p*-cresol is tyrosine metabolism by *Clostridium* species (51), although the specific strains responsible remain to be elucidated. We also detected a DEHP-dependent increase in the level of a metabolite putatively identified as isatin. A recent study in rats found that surgical delivery of indole to the cecum leads to isatin and oxindole accumulation in the brain, while decreasing motor activity (52). Taken together, these results suggest that the gut microbiota could be the source of potentially harmful metabolites previous studies have associated with neurodevelopmental disorders such as ASD.

In addition to the above discussed metabolites, the untargeted analysis detected a large number of compounds whose amounts in the culture were significantly altered by DEHP in a dose-dependent fashion. Only a small fraction (79/1937) of these compounds could be assigned a putative identity, as many of the detected features’ MS/MS spectra could not be matched to available databases for putative identification and subsequent confirmation. The total annotation rate (204/5408) achieved in the present study is comparable to previous metabolomics studies on the gut microbiota, which report annotation rates ranging from 2 (53) to 5% (54). Metabolite identification clearly remains a bottleneck in untargeted metabolomics, and further efforts are warranted to expand coverage of metabolites from commensal gut bacteria in spectral libraries.

The findings of the present study provide evidence that significant alterations could occur even in developed microbiota in response to environmental chemical exposure, and that these alterations include overproduction of selected bacterial metabolites. Several of these metabolites have been found at elevated levels in urine or plasma of subjects diagnosed with neurodevelopmental disorders, in particular ASD. Taken together with recent reports linking phthalate exposure and ASD, our findings suggest the intriguing possibility that the chemical could selectively modify the intestinal microbiota to promote the production of potentially toxic metabolites such as *p*-cresol. Whether metabolites such as *p*-cresol casually contribute to neurodevelopmental disorders or merely indicate dysbiosis associated with these disorders remains to be elucidated. Further work is warranted to determine whether earlier (e.g., immediately after birth) and longer term DEHP exposure would lead to more severe dysbiosis and affect behavioral outcomes.

## Materials and Methods

### Materials

All chemicals were purchased from Sigma Aldrich (St. Louis, MO) unless otherwise specified. DEHP and MEHP were purchased from AccuStandard (New Haven, CT).

### DEHP exposure in mice

Female C57BL/6J mice aged 4-5 weeks were purchased from Jackson Laboratories (Bar Harbor, ME) and maintained on an *ad libitum* chow diet (8604 Teklad Rodent diet, Envigo, Madison, WI). Mice were acclimatized to the animal facility for one week, and were then were given either vehicle (corn oil) or a low or high dose of DEHP (1 or 10 mg/kg bodyweight/day) via oral gavage. Mice were gavaged with DEHP every other day. Fecal pellets were collected immediately before the first gavage (day 0) and on days 7 and 14, flash frozen in liquid nitrogen, and stored at −80°C. On day 14, animals were euthanized using asphyxiation with CO_2_. Animals were handled in accordance with the Texas A&M University Health Sciences Center Institutional Animal Care and Use Committee guidelines under an approved animal use protocol (AUP IACUC 2017-0145).

### In vitro culture of cecal luminal contents

Whole cecum from female C57BL/6J mice (6-8 weeks of age) were harvested and transported to an anaerobic chamber (Coy Lab, Grass Lake, MI) in an anaerobic transport medium (Anaerobe Systems, Morgan Hill, CA). Luminal contents were isolated from the cecum inside the chamber, and then suspended into a slurry in 1 ml of pre-reduced PBS containing 0.1% cysteine by vortexing the suspension for 2 min. Gut microbiota medium (GMM) was prepared as described previously (27). Each batch of cecal luminal contents slurry from a mouse was inoculated into a separate glass test tube containing 10 ml of GMM or GMM supplemented with a low or high dose of DEHP (10 or 100 µM). The inoculated tubes were incubated at 37°C for up to seven days under anaerobic conditions. Tubes containing GMM but without inoculation and incubated under the same conditions were used as negative controls. Culture (or medium) samples were collected on days 1 and 7 post inoculation by removing test tubes from the incubator and centrifuging at 13,000g for 10 min at 4°C. The cell pellet and supernatant were stored at −80°C for further analysis.

### Extraction of metabolites

The fecal pellets and *in vitro* cecal luminal culture samples were homogenized using lysing matrix E beads MO BIO, Carlsbad, CA) on a bead beater (VWR, Radnor, PA) with equal volume of cold methanol and half volume of chloroform. The samples were homogenized for one min on the bead beater, cooled on ice for one minute, and homogenized again for another 2 min. The samples were then centrifuged at 10,000g at 4 °C for 10 min. The supernatant was filtered through a 70-μm sterile nylon cell strainer into a clean sample tube and mixed with 0.6 ml of ice-cold water using a vortex mixer. This mixture was centrifuged again at 10,000g for 5 min to obtain phase separation. The upper and lower phase were separately collected using a syringe while taking care not to disturb the interface. The upper phase was dried to a pellet using a vacufuge (Eppendorf, Hauppauge, NY), and stored at −80 °C until further analysis. Prior to LC-MS analysis, the dried samples were reconstituted in 50 μl of methanol/water (1:1, v/v).

### Untargeted metabolomics

The extracted samples were analyzed for global metabolite profiles using information-dependent acquisition (IDA) experiments performed on a triple-quadrupole time-of-flight instrument (5600+, AB Sciex) coupled to a binary pump HPLC system (1260 Infinity, Agilent). Each sample was analyzed twice, using two different combinations of LC methods and ionization modes to obtain broad coverage of metabolites having varying polarities and isoelectric points (see Supplementary Methods). Raw data were processed in MarkerView (v. 1.2, AB Sciex) to determine the ion peaks. The peaks were aligned based on m/z and retention time (RT), (30 ppm and 2.5 min tolerance, respectively), and then filtered based on intensity (100 cps threshold) to eliminate low quality peaks. An additional filter was applied to retain only monoisotopic ions. The retained ions were organized into a feature table, with each feature specified by m/z and RT. In the case a precursor ion detected by the TOF survey scan triggered an MS/MS scan, the corresponding MS/MS spectrum was extracted from the product ion scan data and added to the feature table. Each feature was searched against spectral libraries in METLIN (55), HMDB (56), and NIST (57). The MS/MS spectrum of each feature was also analyzed using *in silico* fragmentation tools MetFrag (58) and CFM-ID (59). These analyses identified several annotations for many of the features. To systematically determine the most likely identities for these features in the context of murine cecal microbiota metabolism, we applied an automated annotation procedure (‘BioCAn’) that combines the outputs from the database searches and fragmentation analyses with a metabolic model (see below) for the biological system of interest (60). Briefly, BioCAn maps each unique mass in the feature table onto a metabolic network representing the enzymatic reactions possible in the system of interest, and evaluates the likelihood a correct mapping between a detected mass and a metabolite in the network has occurred based on how many other metabolites in the neighborhood of the metabolite in question also map to a detected mass. Features annotated with high confidence were further inspected manually and confirmed by matching the product ion spectrum to standards in the aforementioned reference databases. In cases where no reference MS/MS spectra were available, a pure standard was run to confirm the metabolite identity. Relative amounts of metabolites were quantified using MultiQuant 2.1 (AB Sciex) by manually integrating the corresponding peak areas in the extracted ion chromatograms (XICs).

### Targeted analysis of MEHP

The fate of DEHP in the cecal culture was characterized by quantifying the amount of its major metabolic product, MEHP. Targeted analysis of MEHP utilized a product ion scan experiment as described previously (61).

### 16S rRNA sequencing analysis

Fecal and *in vitro* cecal luminal culture pellets were homogenized and microbial DNA was extracted from the homogenate using the standard protocol for Power soil DNA extraction kit (MO BIO). The V4 region of 16S rRNA was sequenced on a MiSeq Illumina platform using protocols for paired-end sequencing from Kozhich et al. (2013) at the Microbial Analysis, Resources, and Services (MARS) core facility at the University of Connecticut. Sequence reads were quality filtered, denoised, joined, chimera filtered, aligned and classified using QIIME (62, 63). The SILVA database (28) was used for alignment and classification (97% similarity) of the OTUs. The OTU counts were normalized by subsampling to the lowest number of OTUs found in the sample.

### Metabolic model

The OTU tables from QIIME analysis were used to build a metabolic model linking bacterial groups detected in the cecal cultures to metabolites that can be produced by these groups. To select species for inclusion in the model, we tabulated the most abundant OTUs detected in all samples from both days 1 and 7 of GMM culture, with a 0.01% cutoff for relative abundance. The genera associated with these OTUs were searched against the KEGG Organisms database to compile a list of organisms that have a complete genome sequence and an assigned KEGG organism code (64). This list was then manually curated to remove species unlikely to be present in murine cecum (e.g., soil dwelling bacteria and extremophiles) by searching a microbiome database (65) and carefully examining the published literature. From this curated list, we generated a matrix linking an organism to reactions encoded by its genome. First, the KEGG Orthology identifiers (K numbers) and Enzyme Commission (EC) numbers associated with the organism codes were collected using the KEGG REST API. These K and EC numbers were then linked to KEGG reaction identifiers (R numbers). The linkages between organism codes and R numbers were arranged into an organism-reaction (**OR_KEGG_**) matrix, where each element (*i, j*) denotes the presence (‘1’) or absence (‘0’) of a reaction *i* in organism *j*.

The organisms in **OR_KEGG_** accounted for 48 of the 119 most abundant genera in the cecal cultures. The remaining 71 genera were searched against the UniProt database to determine if high-quality genome sequences with functional annotation were available for any of the member strains. After removing species that are unlikely present in the murine cecum, organisms with high-quality functional annotations were added to an organism-enzyme matrix (**OE_UniProt_**), where each element (*i, j*) denotes denotes the presence (‘1’) or absence (‘0’) of an enzyme *i* in organism *j*. The amino acid sequences from each of the remaining organisms lacking annotated genomes were downloaded from GenBank and assigned K numbers using BlastKOALA (66). The resulting linkages between organisms and K numbers were arranged into an organism-orthology matrix (**OK_UniProt_**). The K and EC numbers of these two matrices were linked to R numbers to generate a second organism-reaction matrix (**OR_UniProt_**). The two matrices **OR_KEGG_** and **OR_UniProt_** were combined to produce a final organism-reaction matrix (**OR**) for all detected genera with member species that have high-quality genome sequences. The metabolites associated with each organism were found by linking the reactions with their primary substrate-product pairs as defined by KEGG’s RCLASS data.

### Statistical analysis

OTUs observed only once across all samples were filtered prior to PCA and PLS-DA in MATLAB (v. R2018a). Linear discriminant analysis of the effect size (LEfSe) was used to characterize differences in the OTU counts between samples (67). Effects were considered statistically significant if they were assigned a q-value less than 0.05. A two-tailed t-test with a cutoff *p*-value of 0.05 was used to test for statistical significance of differences in metabolite levels between treatment groups. Pearson correlation coefficients (PCCs) were calculated between OTU counts (relative abundance) and peak areas of metabolites. Statistical significance of the PCCs was determined based on p-values calculated using a two-tailed *t*-test, and corrected for false-discovery rate using the Benjamini-Hochberg (B-H) method (68). Statistically significant correlations (B-H adjusted p-value < 0.05) between OTUs (at the level of genus) and metabolites were visualized in Cytoscape (v. 3.0).

## Acknowledgments

This work was supported by grants from the NSF (1264502) and NIGMS (GM106251) to KL and AJ, the Nesbitt Chair to AJ, and grants from NSF (1337760) and NCI (CA211839) to KL.

## Author Contributions

S.M., R.M., K.L., and A.J. designed the study. M.L., R.M., S.M., C.H., and R.C.A. performed the experiments. M.L., R.M., and N.A. analyzed the data. M.L., R.M., K.L., and A.J. wrote the manuscript.

^1^M.L. and R.M. contributed equally to this work.

